# Decrypting the microbiota on the black soldier fly’s (*Hermetia illucens* L., Diptera: Stratiomyidae) egg surface and their origin during development

**DOI:** 10.1101/2022.12.22.520758

**Authors:** Carina D. Heussler, Thomas Klammsteiner, Katharina T. Stonig, Heribert Insam, Birgit C. Schlick-Steiner, Florian M. Steiner

**Author notes:** These authors contributed equally as first authors (Carina D. Heussler, Thomas Klammsteiner). These authors contributed equally as senior authors (Birgit C. Schlick-Steiner, Florian M. Steiner). Corresponding author: Carina Desirée Heussler.

## Abstract

The increasing global population leads to a soaring demand for protein for food and feed, and also challenges in organic waste management are growing. All this leads to environmental stress, causing biodiversity loss and an increase in greenhouse gas emissions. Alternative and sustainable animal protein sources are needed to reduce the negative environmental impact of food production. In the last years, the black soldier fly (BSF) has been proposed to substitute animal protein, since BSF may consume and reduce a variety of waste organic matter. Successful industrial rearing of BSF depends on a flourishing reproduction of adults, which is influenced not only by environmental but also by physiological factors. The BSF female oviposits single eggs into clutches close to decomposing organic matter and conspecific eggs. Studies have shown that microbes play a significant role in the oviposition of BSF eggs. In this study, we focus on the surface microbiota of the egg and its origin. We investigated if the microbiota is inoculated before, during, or actively after oviposition. For this purpose, we analysed the microbiota in the larval haemolymph and the gut of larvae raised on sterilized and non-sterilized feed, the pupal cell pulp, the wash of the egg-laying apparatus and the eggs directly collected after oviposition, the ovarian eggs and the empty female abdomen, the eggs with contact to adult BSF, and sterilized eggs. The bacterial communities were identified through 16S rRNA gene sequencing to assess the stage in BSF development during which the microbial colonization of the egg surface occurs. We demonstrated that bacteria differ among life stages resulting in a shift during BSF development from dominance of *Enterobacteriaceae* during the larval stage to dominance of *Burkholderiaceae spp*. in all analysed eggs. A predominant microbiota is present before oviposition and persists through all life stages, however, the overall population’s structure successively shifts during development. A better understanding of egg surface microbiota and oviposition attractants could significantly increase egg production and facilitate the mass harvesting of BSF larvae.

**Graphical abstract.**
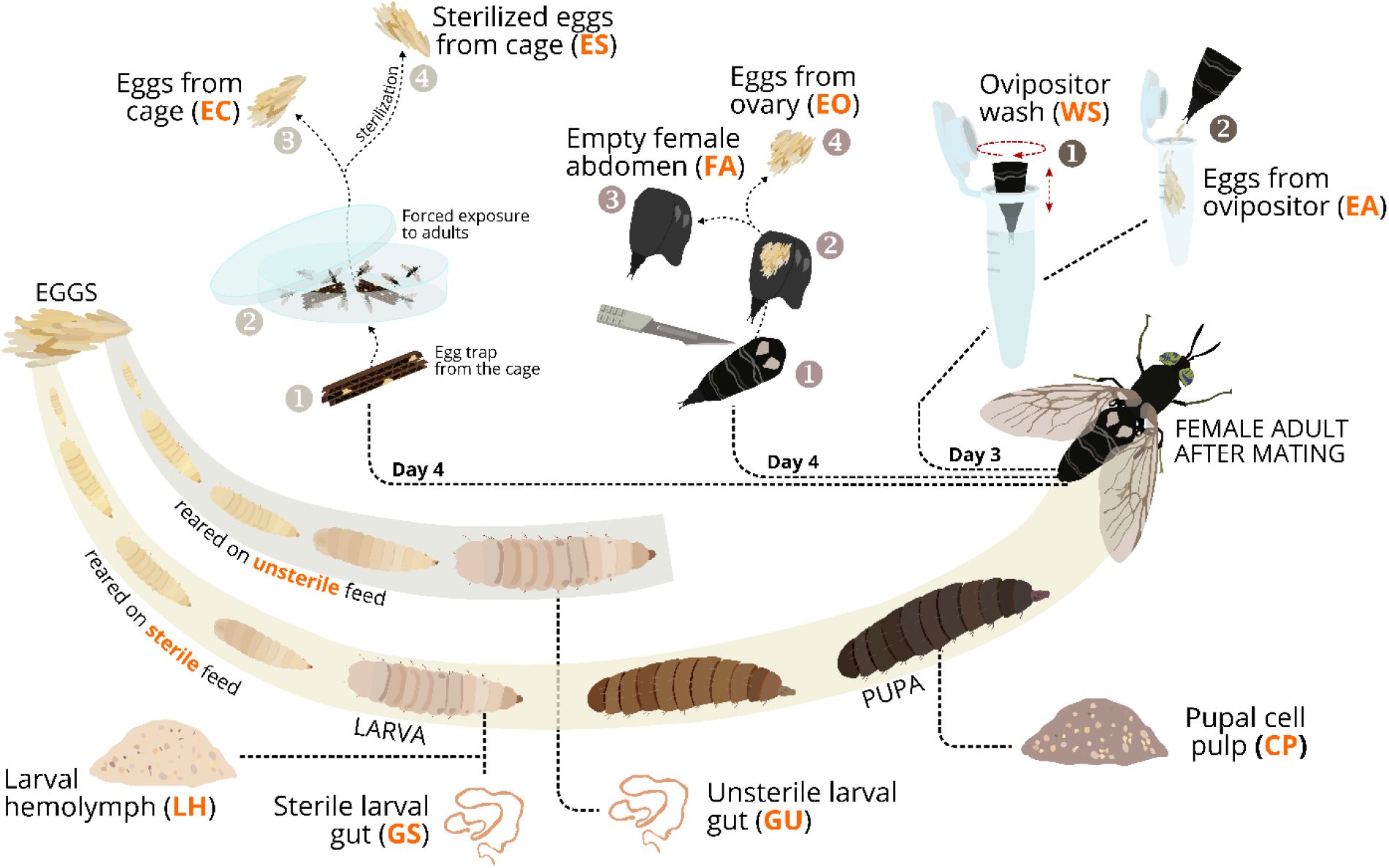
Illustration of the sampling procedure to assess egg surface microbiome of black soldier fly and its origin: Larval fed with unsterile (GU) and sterile (GS) feed and the larval haemolymph of GS (LH), the pupal cell pulp (CP), and from the female adults after mating, a wash of the egg-laying apparatus (WS) and the afterwards placed eggs of the egg-laying apparatus (EA), eggs collected from the ovary (EO) and the empty female abdomen (FA), eggs collected from a fly cage after forced exposure to adults (EC) and sterilized (ES).

## Introduction

The rapid population growth, accelerating urbanization, and rising incomes lead to increased demand for food and feed and to a growing challenge in managing organic wastes (Alexandratos and Bruinsma, 2012; Rosegrant and Msangi, 2011; Van Huis and Oonincx, 2017). In turn, this leads to an intensification of environmental stress through deforestation, overfishing of the oceans, and greenhouse gas emissions on account of the production of animal proteins such as beef, pork, and chicken (Calicioglu *et al*., 2019). To meet the demands of animal protein resources, meat production is estimated to increase by more than 75% until 2050, and global fish production is projected to be 30 million tons higher in 2030 compared with 2016 (Alexandratos and Bruinsma, 2012; Van Huis and Oonincx, 2017). Furthermore, the demand for animal and/or plant-based protein-rich animal feed like fish- or soybean meal will surge.

In 2021, 20% of wild-caught fish was processed into fishmeal for aquaculture purposes (Thrastardottir *et al*., 2021). The current food system is responsible for 80% of deforestation and 29% of greenhouse gas (GHG) emissions. Additionally, agriculture uses 34% of all land globally and withdraws 70% of the freshwater used globally for watering. As a result, the current food system is estimated to be responsible for 68% of biodiversity loss, which effect is predicted to grow even further (Nations, 2021; Thrastardottir *et al*., 2021). Moreover, about 1.7 billion tons of all food produced is wasted (Surendra *et al*., 2020). The resulting organic waste is mostly not properly managed and has a significant negative impact on the environment (3.3 billion tons of GHG emissions, per year) and the economy (total cost of 1.2 trillion dollars per year) (FAO, 2014; Surendra *et al*., 2020; Zhou *et al*., 2022).

To reduce the environmental impact of food production, alternative sources of animal protein are needed. Insect biomass as an alternative animal protein source has been reported to have a lower environmental impact than other sources of animal protein (Guiné *et al*., 2021). Insect farming emits less GHG, uses less land, and requires lower inputs of feed and water due to higher conversion efficiencies (Guiné *et al*., 2021; Matos *et al*., 2021).

In the last years, the black soldier fly (*Hermetia illucens*, L., BSF) has been identified as a key player to substitute animal protein for food and feed (Smetana *et al*., 2019; Wang and Shelomi, 2017). The BSF adults have a reduced digestive tract; they primarily survive on reserves accumulated during the larval stage. Hence, BSF larvae (BSFL) are high in protein (40-44% dry matter base) and fat (up to 49% dry matter base) and contain several micro-and macronutrients important for livestock health and development (Makkar *et al*., 2014). The larvae are able to digest a large variety of organic matter such as food waste, faecal sludge, manure, and agro-industrial by-products (Wang and Shelomi, 2017). The undigested residues mixed with BSFL excrements find use as organic fertilizer, capable to substitute or replace mineral fertilizers (Klammsteiner *et al*., 2020a). By converting organic wastes into nutrient-rich insect biomass suitable for feedstock production with organic fertilizer as the main process by-product, the BSF can contribute to circular economy goals (Liu *et al*., 2022; Walter *et al*., 2020). It even has been reported that BSFL can reduce the methane production of swine manure by 86% and that the direct GHG emissions are lower than conventional composting (Matos *et al*., 2021; Mertenat *et al*., 2019).

For these and many more reasons, the BSF is an ideal candidate for the industrialization of insect farming (Miranda *et al*., 2020). Successful large-scale rearing of BSF depends on a flourishing reproduction of adults (Heussler *et al*., 2022). The reproductive performance of the BSF is influenced by physiological and environmental factors as well as technological parameters (Boaru *et al*., 2019). The female adults oviposit single eggs in clutches (up to 900 eggs) close to decomposing organic matter and are attracted by conspecific eggs (Nakamura *et al*., 2016). Zheng *et al*. (2013b) showed that bacteria isolated from conspecific eggs attract gravid females, presumably by emissions of volatile organic compounds. A better understanding of the interkingdom communication between microbes and BSF females, and in particular regulation of oviposition, can significantly increase the egg production and mass harvesting of larvae (Boaru *et al*., 2019).

In this study, we focused on the bacterial communities on the egg surface and their origin. We hypothesized that the microbiota on the egg surface will either be inoculated before oviposition, during oviposition, or actively after oviposition by adult flies. We sampled the microbiota in ten approaches, that is, from the larval haemolymph (LH) and the larval guts fed with sterilized (GS) and non-sterilized feed (GU), the pupal cell pulp (CP), the wash of the egg-laying apparatus (WS) and the eggs directly collected after oviposition (EA); the ovarian eggs (EO) and the empty female abdomen (FA); eggs of the fly cage with contact to adult BSF (EC), and sterilized eggs (ES). The microbiota were identified through 16S rRNA gene sequencing to assess the stage in BSF development when the microbial colonization of egg surface occurs.

## Material and Methods

### Breeding of black soldier flies

The BSF colony was reared in an ICH750eco climate chamber (Memmert, Schwabach, Germany) at a temperature of 27 °C (± 0.5 °C) and 60% relative humidity and a 16:8 h (L:D) photoperiod (Tomberlin *et al*., 2009). BSFL were kept in black polypropylene boxes (180 × 120 × 80 mm) sealed with plastic lids with integrated nets to allow aeration. Larvae were fed twice a week *ad libitum* with a 40:60 (w/v) mixture of ground chicken feed (Grünes LegeKorn Premium, Landwirtschaftliche Genossenschaft, Klagenfurt, Austria) and tap water until prepupation (Sheppard *et al*., 2002). When the transition to the pupal stage started, migrating prepupae were guided into collection boxes via a ramp installed on the inner side and a pipe attached on the outer side of the larva boxes, thereby allowing their self-harvesting. Thereafter, pupae were kept in white plastic cups (50 ml) and covered with wood shavings (Dehner Terra, Rain, Germany). After eclosion, flies were collected manually into transparent polypropylene cages (390 × 280 × 280 mm) with a net (fibreglass; 200 × 300 mm, mesh size 2 × 2 mm) for aeration integrated into the lid of the cage. The fly cage was illuminated with light-emitting diodes panels (Y51515227 184210, Barthelme, Nuremberg, Germany) in a 16:8 (L:D) photoperiod (Heussler *et al*., 2018). On two opposite walls of the fly cage, one piece of corrugated black polypropylene cardboard each (henceforth termed flutes) was installed for oviposition by using magnets. A glass test tube filled with tap water and plugged with a piece of cellulose paper was provided as a water source. On Day 4 after manually transferring the flies into the cages, the eggs were harvested from inside the flutes.

### Experimental setup and sampling

A flute containing ten egg clutches was transferred into larva boxes, placed above chicken feed freshly mixed with water, and held in place using toothpicks. To prevent larvae from escaping, dry ground chicken feed was spread along the inner edges of the box as described in Addeo *et al*. (2022). To avoid the introduction of exogenous microorganisms into the feed, the larvae used in this study were fed with ground, autoclaved chicken feed mixed 60% (w/v) with distilled water *ad libitum* every other day. Ten distinct sampling approaches were applied to assess when, where, and which bacterial and fungal communities occurred during BSF development, and to determine the source of microbial colonization. Additionally, a control treatment with larvae fed with unsterilized chicken feed was included. Samples for subsequent DNA extraction were collected in triplicates; for each replicate, five individuals were pooled, except for WS, where ten individuals were pooled due to low amounts of DNA; for details, see section “DNA extraction”.

### Collection of larval haemolymph (LH) and guts for non-from sterilized (GU) and sterilized diet (GS)

Nine days after hatching (DAH), about 50 larvae were collected and stored at −20 °C, in estimation of half-life at 18 days until first prepupation through self-observation and literature (De Smet *et al*., 2018). Frozen larvae were thawed for two min in a solution of 50:50 5% bleach (Danklorix, CP GABA, Hamburg, Germany) and Milli-Q (Merck, Darmstadt, Germany) water for surface sterilization (Amiri *et al*., 2018; Hausmann *et al*., 2021) and then placed in a 50-ml tube filled with pure Milli-Q water. To assure complete extraction, the gut was extracted by pulling out the anus using sterile forceps and then transferring it into a sterile microcentrifuge tube (at least 0.05 g/replicate). The remaining haemolymph was placed in an empty and sterile microcentrifuge tube (at least 0.10 g/replicate). The same procedure was used to collect the gut of the larvae fed with the non-sterilized diet.

### Collection of pupal cell pulp (CP)

About 50 pupae were collected 19 DAH and stored at −20 °C. Frozen pupae were thawed for two min in a solution of 50:50 5% bleach and Milli-Q water for surface sterilization, and afterwards, pupae were placed in a 50 ml tube filled with Milli-Q water. Each pupa was cut along both lateral sides with sterile scissors and the cell pulp was scraped out with a sterile spatula. All five pupa replicates were collected into an empty and sterile microcentrifuge tube (at least 0.10 g/ replicate).

### Collection of wash of the egg-laying apparatus (WS) and eggs immediately after oviposition (EA)

To assure easy handling and to prevent flies from escaping while collecting gravid females, the number of flies per cage was limited to a density of 0.0033 flies/cm^3^. 50 females and 50 males aged 24 h were released into the fly cages. A total of three fly cages were used to obtain enough females for all sampling approaches, and sampling of individuals occurred randomly across all cages. The fly cages were kept under the same conditions (see section “Breeding of black soldier flies”). To obtain enough gravid females and assuming that females start oviposition on Day 4 after transferring flies to the cages (Heussler *et al*., 2018), the collection of females for this treatment was set on Day 3 after transferring flies to the cages. Gravid females were collected manually one at a time. Each female was held above a sterile microcentrifuge tube filled with 700 μl lysis buffer SL1 (NucleoSpin Soil kit, Macherey-Nagel, Düren, Germany), and the ovipositor was dipped into the liquid and moved in circles for one min to wash microbes off of the ovipositor’s surface (WS). Thereafter, the female was decapitated to induce oviposition. Each female was held above a sterile microcentrifuge tube to allow oviposition into the tube (EA; at least 0.05 g/replicate).

### Collection of the ovarian eggs (EO) and the empty female abdomen (FA)

Approximately 20 gravid females were collected on Day 4 after transferring flies to the fly cages and stored at −20 °C. Frozen females were thawed for two min in a 50:50 solution of 5% bleach and Milli-Q water for surface sterilization and placed in a 50 ml tube filled with Milli-Q. To access the ovary, each female was cut along both lateral sides of the abdomen with sterile scissors, and the ovary was collected into a sterile microcentrifuge tube using a sterile spatula (at least 0.05 g/replicate). The remaining abdomen of the female was separated from the thorax and collected into a sterile microcentrifuge tube (at least 0.05 g/replicate).

### Collection of eggs from the fly cage after contact with adult BSF (EC)

The remaining flies in the fly cages were allowed to oviposit. On Day 6 after transferring the flies into the fly cage, the flutes were collected into a sterile petri dish. Five females and five males were introduced into the petri dish and left for one hour at 27 °C and 60% RH. This procedure assured contact between the eggs and adults, and enabled inoculation of eggs with adult-derived microbes. Thereafter, five egg clutches were collected into sterile microcentrifuge tubes (at least 0.05 g/replicate).

### Collecting of sterilized eggs (ES)

Five egg clutches were collected into sterile microcentrifuge tubes (at least 0.05 g/replicate). The tubes were filled with 700 μl of a 50:50 mixture of 5% bleach and Milli-Q Water, vortexed for 10 s, and incubated for 2 min. The tubes were centrifuged (30 s at 11,000 × g), and the supernatant was removed. The pellet was washed following these steps: 700 μl of Milli-Q water was added; vortexed for 10 s; centrifuged (1 min /11,000 × g), the supernatant was removed, and these steps were repeated at least five times until the smell of bleach was no longer noticeable but before the egg surface started to break.

### Viability of sampled eggs

To assess the viability of the eggs after the sampling procedures, additionally, three egg clutches were collected for each of the sampling procedures and placed in a sterile Eppendorf tube above non-sterilized feed and controlled daily for hatching larvae.

### Differences in larval growth on sterilized and non-sterilized feed

To assess whether there are differences in larval growth reared on sterilized and non-sterilized feed, we performed a separate feeding trial. BSFL were reared in triplicates (90 larvae per replicate) on sterilized and non-sterilized chicken feed 40:60 (w/v) in 90 x 90 x 40 mm boxes covered with cellulose paper. The BSFL were fed a fresh weight of 100 mg/larva/day. Every other day, three times ten larvae (without replacement) were randomly selected from each box and weighted to track their biomass gain.

### DNA extraction

DNA was extracted using the NucleoSpin Soil kit (Macherey-Nagel, Düren, Germany) following the manufacturer’s protocol with some modifications: the lysed sample of the larval haemolymph was vortexed for 10 min, and the other samples for 5 min. Prior to precipitation with SL3 Buffer, the supernatant was moved to a new collection tube. DNA was eluted twice using 20 μl SE Elution Buffer each. DNA concentration and quality were checked via gel-electrophoresis and UV-Vis spectrophotometry (NanoDrop 1000c, Thermo Fisher Scientific, Waltham, MA, USA). DNA was stored at −20 °C until further processing.

### 16S rRNA gene amplicon sequencing

The UV-Vis spectrophotometry showed low DNA concentrations for some samples. Therefore, an enrichment PCR of all samples was performed by the sequencing provider to amplify the DNA. The enrichment was performed by diluting (2.5 ng/μl) the samples, followed by an enrichment PCR with locus-specific primers (V34:F TCGTCGGCAGCGTCAGATGTGTATAAGAGACAGNNNNNCCTACGGGNGGCWGCAG; R GTCTCGTGGGCTCGGAGATGTGTATAAGAGACAGNNNNNGACTACHVGGGTATCTAATCC, red = locus-specific sequences). Then, a 1^st^-step PCR with locus-specific primer and Illumina overhang and a cleanbead purification were performed, followed by a 2^nd^-step PCR with index primer and another cleanbead purification. The final libraries were pooled, and a final cleanbead purification of the pool was carried out. Illumina MiSeq amplicon 16S genetic sequencing was performed by Microsynth AG (Balgach, Switzerland) using a 2 × 250 bp paired-end approach with the universal bacterial primers 341f (5’-CCTACGGGRSGCAGCAG-3’) and 802r (5’-TACNVGGGTATCTAATCC-3’) targeting the V3-4 region on the 16S rRNA gene. Library preparation was performed by the sequencing provider based on a Nextera two-step PCR including purification, quantification, and equimolar pooling. In addition, the ITS2 genetic region was sequenced using the ITS3f (5’-GCATCGATGAAGAACGCAGC-3’) and ITS4r (5’-TCCTCCGCTTATTGATATGC-3’) primer pair. However, due to the low quality of the reads resulting from this sequencing job, we decided to exclude the data on fungal communities from further analyses and interpretation.

### Processing and analysis of sequencing data

Raw reads generated by targeting the V3-V4 genetic region were filtered, trimmed, and dereplicated in DADA2 v1.8 (Callahan *et al*., 2016). After inferring amplicon sequence variants (ASVs), the paired forward and reverse reads were merged, and chimeras were removed. Taxonomy was assigned using the reference databases SILVA (v.132; Quast *et al*. (2012)). The data were visualized using ampvis2 (v.2.7.4; Andersen *et al*. (2018)) and ggplot2 (v.3.3.5; Wickham (2016)). Venn diagrams were created using the MicEco package (v.0.9.17; Russel (2022)). Reproducible documentation of sequence processing and data analysis as well as download options for relevant data can be accessed via https://tklammsteiner.github.io/eggsurfacemicrobiome.

### Statistical analysis

Alpha diversity (Shannon index) and linear discriminant analysis of effect size (LefSe; threshold at LDA score (log10) >= 2) were calculated using the microbiome (v.1.16.0; Lahti *et al*. (2017) and microbiomeMaker (v. 1.0.1; Yang (2021) package, respectively. Differences between means of alpha diversity indices were calculated via Wilcoxon test. Permutational analysis of variance (PERMANOVA) was calculated based on Bray-Curtis dissimilarity values using the anosim function (permutations = 1000) in vegan (v.2.5-7; Oksanen *et al*. (2020)). Pairwise differences in microbial community composition of treatment groups were assessed using the pairwise.perm.manova function (nperm = 1000) with subsequent Bonferroni correction in RVAideMemoire package (v.0.9.81; Hervé (2022)). The results were considered statistically significant when their p-value was < 0.05.

## Results

### An extensive shift in family-level relative abundance during BSF development

An average of 31,698 ± 11,232 raw reads per library were generated by Illumina MiSeq amplicon sequencing. After filtering, denoising, and chimera removal, 26,125 ± 8,914 high-quality reads remained, which were further rarefied to the smallest sample size (16,167 reads) before subsequent biostatistical analysis. Due to an inadequately low read number (679 reads), sample EA1 (replicate 1 of eggs from ovipositor) was removed as an outlier in the process of subsampling. The highest relative abundance for bacteria in all larval stages was *Enterobacteriaceae*, though less dominant for LH, followed by *Enterococcaceae* (Fig 1A, Fig S1). In LH, the families *Aerococcaceae, Bacillaceae*, and *Burkholderiaceae* were also highly abundant. In terms of genus-level representatives of the *Enterobacteriaceae, Morganella* sp. was most abundant for all larval stages (GU, GS, and LH), followed by *Escherichia* sp. and *Proteus* sp. (Fig 1B). The abundance of *Enterobacteriaceae* decreased during the prepupal stage and was similarly abundant as *Burkholderiaceae* in the CP samples. Within the family of *Enterobacteriaceae, Providencia* was the most abundant genus in CP. This trend shifted slightly in the adult stage where the abundance of *Enterobacteriaceae* increased again, but *Burkholderiaceae* were still highly present. In all the egg stages, *Burkholderiaceae* accounted for most of the classified sequences, with the group of *Burkholderia-Caballeronia-Paraburkholderia* sp. as the most abundant genera dominating EC and ES samples. Similarities were found in microbiota composition between the FA, WS, and EA samples, showing the presence of *Xanthomonadaceae, Micrococcaceae*, and *Staphylococcaceae*.

**Fig 1.**
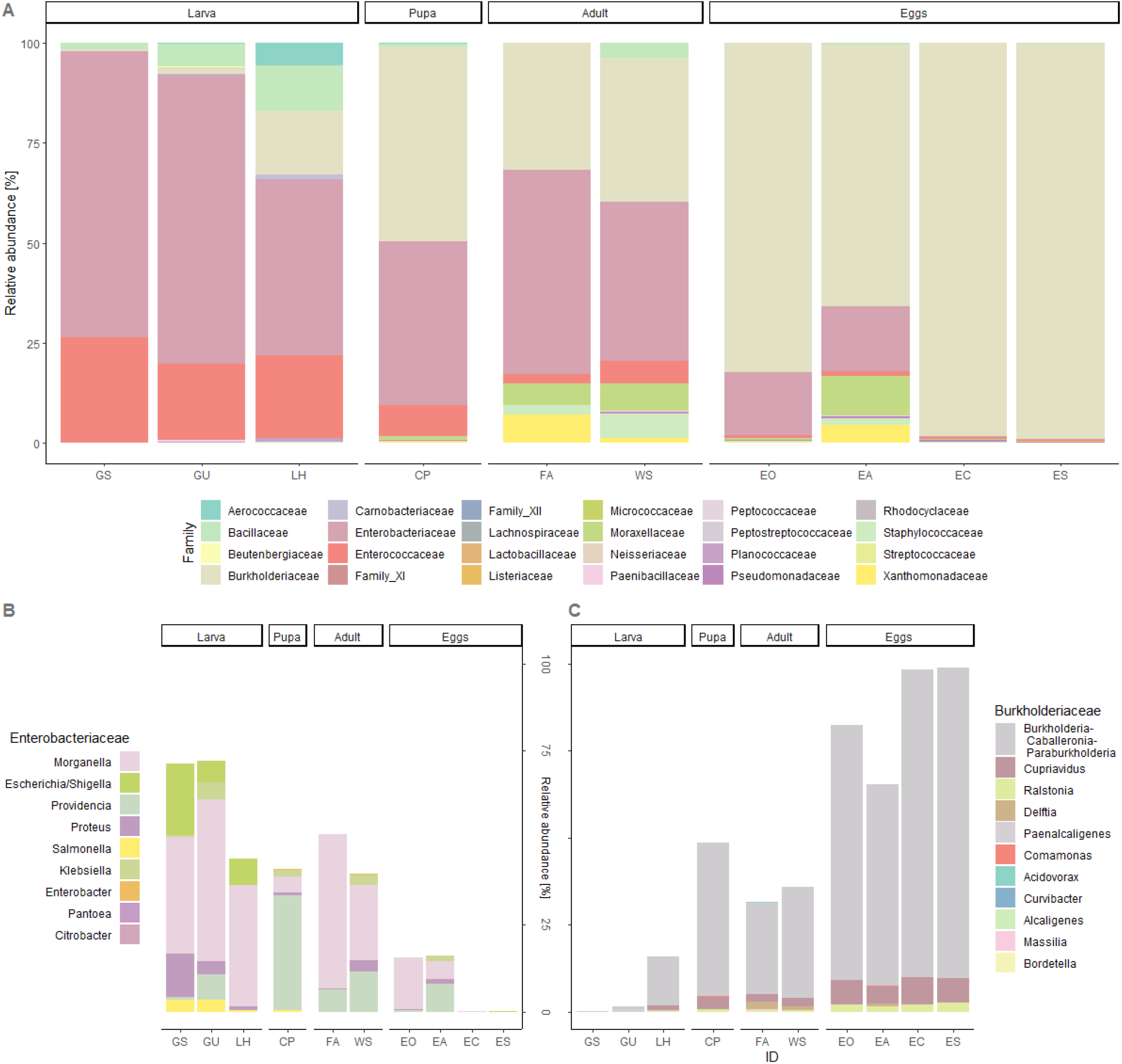
A) Community composition at family level. Genus-level composition of the families B) Enterobacteriaceae and C) Burkholderiaceae in different life stages of the black soldier fly: Larval fed with unsterile (GU) and sterile (GS) feed and the larval haemolymph of GS (LH), the pupal cell pulp (CP), and from the female adults after mating, a wash of the egg-laying apparatus (WS) and the afterwards placed eggs of the egg-laying apparatus (EA), eggs collected from the ovary (EO) and the empty female abdomen (FA), eggs collected from a fly cage after forced exposure to adults (EC) and sterilized (ES).

### Diversity in the egg-surface microbiome during BSF development and unique and shared ASVs

As expressed by the Shannon diversity index (Fig 2), the egg stage had a significantly lower diversity compared with the larval (p = 0.020) and pupal stages (p = 0.038) and a highly significant decrease in diversity compared with the adult stage (p = 0.004). In the larval stage, GU had the highest diversity (H’ = 2.6 ± 0.9) and LH the lowest (H’ = 2.2 ± 1.1). For the adult stage, the WS had a higher diversity (H’ = 2.8 ± 0.3). For the egg stage, the highest diversity was observed in the EA samples (H’ = 2.2 ± 0.8), while all of the ES samples had a relatively low diversity (H’ = 1.1 ± 0.1). The highest distance in the network analysis between communities can be observed from LH and GS to EC and ES (Fig S2).

**Fig 2.**
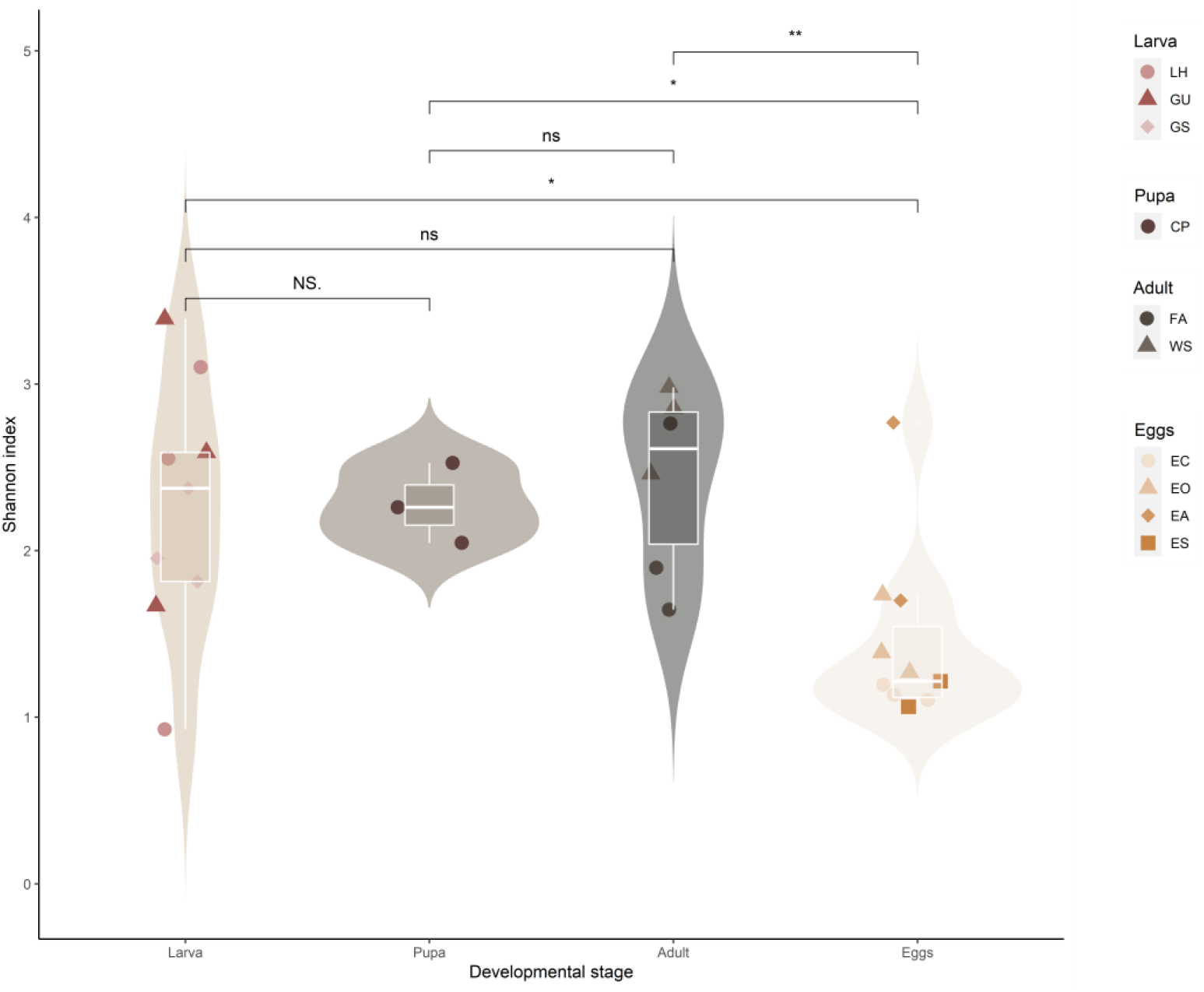
Shannon diversity index (NS: p = 1, ns: p> 0.05, *: p<= 0.05; **: p <= 0.01) for the microbial communities of various black soldier fly life stages: Larval fed with unsterile (GU) and sterile (GS) feed and the larval haemolymph of GS (LH), the pupal cell pulp (CP), and from the female adults after mating, a wash of the egg-laying apparatus (WS) and the afterwards placed eggs of the egg-laying apparatus (EA), eggs collected from the ovary (EO) and the empty female abdomen (FA), eggs collected from a fly cage after forced exposure to adults (EC) and sterilized (ES).

The highest number of unique ASVs (ASVs not shared with other life stages) was found in the larval stage (118 ASVs, Fig 3A), while the lowest number of unique ASVs was found in the pupal stage (30 ASVs). The most shared ASVs were found between the adult and egg stages (28 ASVs) and the lowest shared ASVs were between larva and egg, and pupa and adult (2 ASVs). Among the sampling approaches WS, EA, FA, and EO (Fig 3B), the highest unique ASVs for adults was found via WS (61 ASVs), whilst via FA only 6 unique ASVs were found. Originating from the same treatment, the highest number of ASVs for the egg stage was found in the EA (35 ASVs), and similarly low to FA was the number of ASVs found in EO (7 ASVs). The highest number of shared

**Fig 3.**
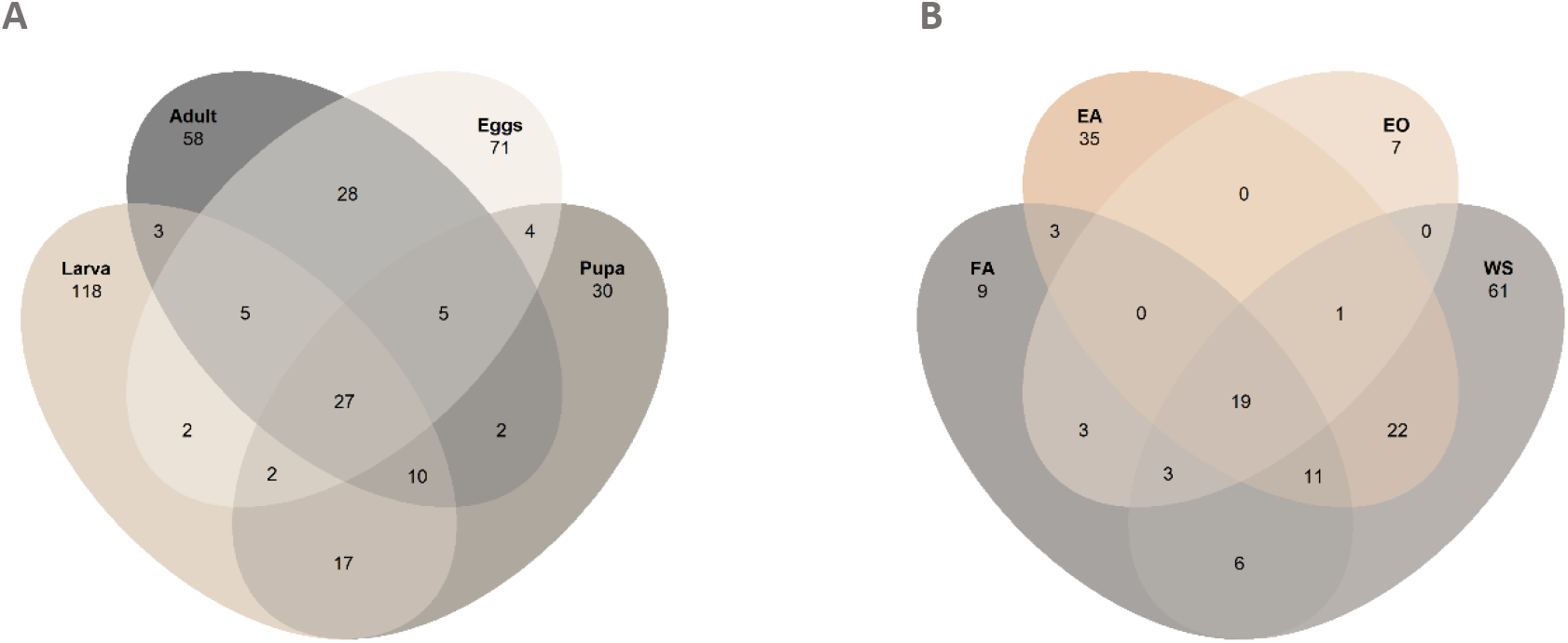
Venn diagram showing unique and shared ASVs for A) different life stages of black soldier fly and B) comparison between adult and egg stages collected from the female adults after mating, a wash of the egg-laying apparatus (WS) and the afterwards placed eggs of the egg-laying apparatus (EA), eggs collected from the ovary (EO) and the empty female abdomen (FA).

ASVs was found between WS and EA (22 ASVs), while FA and EO only shared 3 ASVs. Between EO and WS, and FA and EO, the shared ASVs were 0. However, WS and FA shared 6 ASVs.

Biomarker analysis based on LefSe further distinguished significantly overrepresented bacterial genera for larval, pupal, adult, and egg developmental stages. The *Burkholderia-Caballeronia-Paraburkholderia* group was found characteristic for egg samples, while the genera *Acinetobacter*, *Staphylococcus*, and *Stenotrophomonas* were similarly overabundant in adult samples. *Providencia*, which also showed high relative abundances in adults and eggs, was by far the most overrepresented genus in pupal samples. Among the investigated BSF life stages, the complexity of the larval gut microbiota put forth the most biomarker taxa passing the threshold of LDA >= 2, with *Morganella, Melissococcus*, and *Escherichia/Shigella* scoring highest.

PERMANOVA based on Bray-Curtis distances confirmed that there were significant differences in microbial communities across stages (p = 0.001; larvae, pupae, adult, egg), as well as across tissue samples (p = 0.001; larval guts, larval haemolymph, adult haemolymph, ovipositor, ovarium, and eggs) (Tab S1). Pairwise significant differences were shown in microbial community composition across all developmental stages, except between pupae and adults, and across all tissue samples, except between tissue of ovarium and tissue of haemolymph and ovipositor (Tab S2).

### Viability of eggs and larval growth

Only for the EA and EC sampling procedure, hatching larvae could be observed and showed successful growth (Fig S3). The eggs from the EO and ES sampling procedures were not viable. No significant differences could be found for the larval growth feed with sterilized and non-sterilized feed (Fig S4).

## Discussion

This study aimed to decrypt the microbiota on the surface of the BSF eggs and to assess at which developmental stage and how the egg’s inoculation occurred. Our results indicate that a gradual shift in bacterial community structure occurs during BSF development, from an *Enterobacteriaceae*-dominated community in larval stages to a *Burkholderiaceae*-dominated community in egg stages. Furthermore, the results indicate that innoculation of the dominant bacterial communitiy on the eggs surface occurs before oviposition, as the relative abundace in the EO samples show.

The composition of bacterial communities in the gut of the GU and GS larvae was similar, indicating little to no influence of sterilization of the feed. In studies where BSFL were fed with unsterilized chicken feed, *Gammaproteobacteria* (*Enterobacteriaceae*, *Morganella sp*.) and *Bacilli* (*Enterococcus* sp., *Lactococcus* sp., Fig 1A, 1B and 4) were the most abundant phyla, thus supporting our results (Klammsteiner *et al*., 2020b; Klammsteiner *et al*., 2021; Tegtmeier *et al*., 2021). Genera such as *Morganella* sp., *Enterococcus* sp., and Providencia sp. were significantly overrepresented in larval and pupal samples and have previously been found to play a key role in the BSFL core gut microbiome (Fig. 4; (Klammsteiner *et al*., 2020b)). In LH, the relative abundance of *Enterobacteriaceae* decreased, though still dominant, while *Bacilli* and *Burkholderiaceae* increased. *Bacilli* have often been associated with BSF larvae; they promote their growth by fermenting their food (Klammsteiner *et al*., 2020b; Yang *et al*., 2021; Yu *et al*., 2011), whereas *Burkholderiaceae* have been rarely detected, and their role in the BSF life cycle is unknown. In a study performed by Zheng *et al*. (2013b), *Burkholderiaceae* were identified through all life stages of BSF, though classified as a minor component of the fly’s microbiome. In the CP samples, *Enterobacteriaceae* were almost as abundant as *Burkholderiaceae*. This trend shifted in the adult stage, in which *Enterobacteriaceae* made up the major share of reads, though *Burkholderiaceae* were still heavily represented. In addition, the groups of *Xanthomonadaceae, Micrococcaceae* and *Staphylococcaceae* had an increased presence in the adult stage. These families have previously been associated with BSFL (Klammsteiner *et al*., 2020b; Zheng *et al*., 2013a; Zheng *et al*., 2013b). In other insect species such as the pumpkin fruit fly (*Zeugodacus tau*), the same bacterial families have been reported to possibly serve as a source of vitamins, nitrogen, and amino acids, and some of them can be transmitted vertically from the parents to their offspring (Noman *et al*., 2021). A similar population structure of bacteria was found in the EA treatment, indicating that an inoculation through the egg-laying apparatus occurs, as this treatment consisted of freshly oviposited eggs. Furthermore, the EO samples showed a heavy dominance of *Burkholderiaceae* and a strong presence of *Enterobacteriaceae*, but unlike EA, no other bacterial groups were highly represented, indicating an inoculation at the time of oviposition. Interestingly, most other bacterial groups on the older eggs (EC and ES) were outcompeted by *Burkholderiaceae*. Some members of the *Burkholderiaceae* can function as opportunistic pathogens and some have the capacity to degrade chlororganic pesticides (Zheng *et al*., 2013b). A study has shown that a stinkbug (*Saccharum officinarum*) may harbour some fenitrothion-resistant *Burkholderia* from the environment, and these bacteria can be an easy way for the insect to detoxify itself from insecticides (Minard *et al*., 2013). *Burkholderia-Caballeronia-Paraburkholderia* sp. were the most dominant genera found among the *Burkholderiaceae* throughout all life stages and were identified as characteristic colonizers of BSF eggs (Fig 1C, 4). These taxa can interact with their host by supplementing specifically required forms of nitrogen and other nutrients to ensure normal development of, for example, xylophagous insects (Ge *et al*., 2021). The features of bacterial communities across the life stages of BSF indicate that some important microbes are tightly linked to the fly’s development and growth, which includes maturation of the immune system, resistance to chemical substances (e.g., insecticides), supplementation of nutrients, and digestion (Hu *et al*., 2020).

**Fig 4.**
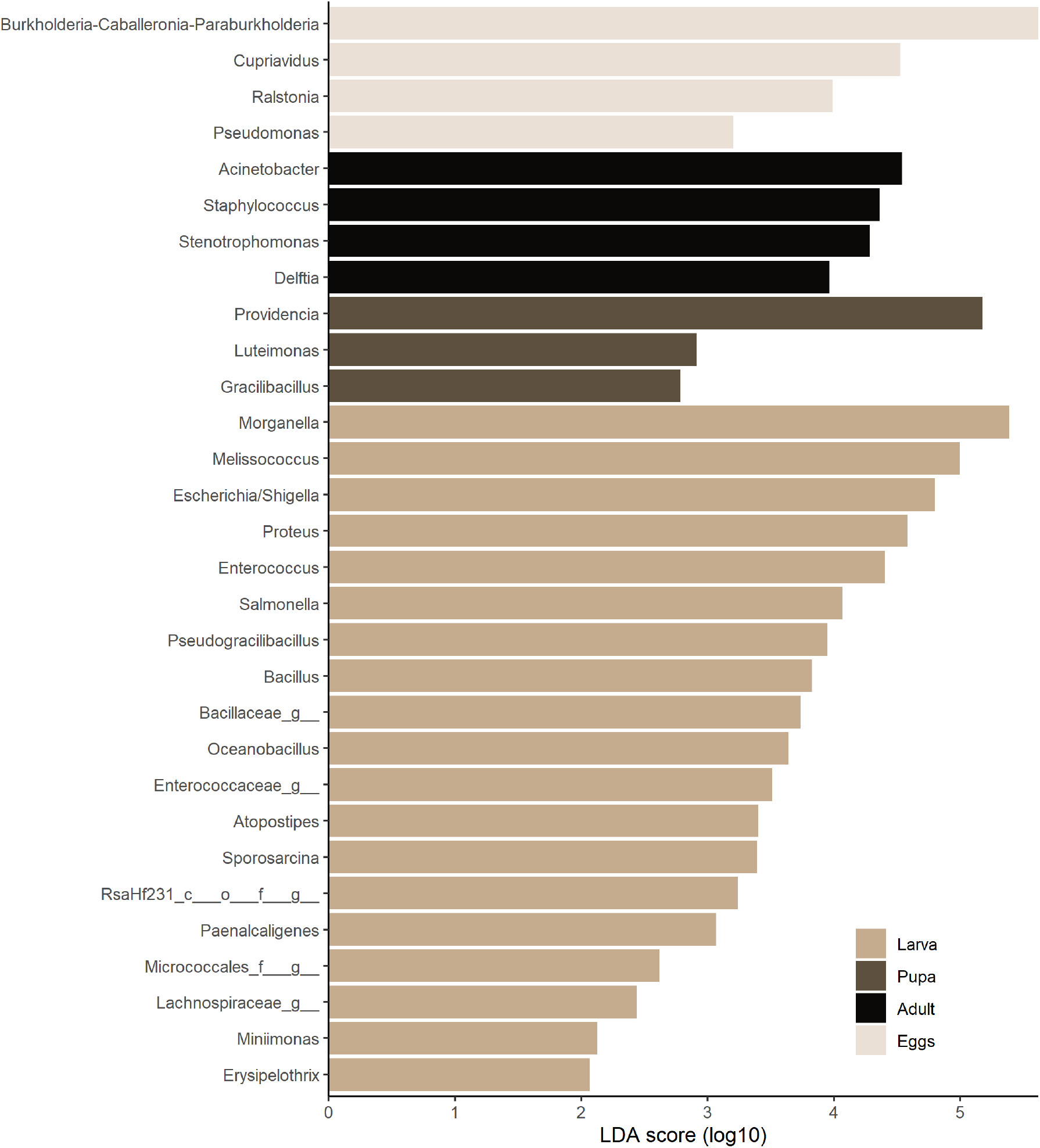
Linear discriminant analysis of effect size (LefSe) identified characteristic bacterial genera for each developmental stage. Samples were grouped based on their life stage, whereas Larva consisted of GU, GS, and LH samples; Pupa consisted of CP samples, Adult consisted of FA and WS samples; Eggs consisted of EO, EA, EC, and ES samples. The threshold for biomarker identification was set to LDA score (log10) >= 2.

Similarly, the Shannon diversity index indicates that bacterial community composition experiences changes in community complexity during the BSF life cycle (Fig 2). The bacterial diversity was higher during larval and adult stages and decreased for the pupal stage and reached a significantly low level for egg stages. This is not surprising, considering BSF eggs are immobile and pupae are semi-immobile and do not feed anymore and, therefore, have less exposure to environmental and transient microbes (Zheng *et al*., 2013b). The lowest diversity was observed for the ES treatment indicating that sterilisation of the egg surface can be considered effective. However, to assess if trans-generational immune priming occurs via bacterial injection into the developing egg, further analyses testing the sterilization should be performed (Freitak *et al*., 2014). Zheng *et al*. (2013a) showed that sterilized eggs lead to a lower percentage of oviposition. This suggests that bacteria of unsterilized eggs attract conspecific gravid females known for other dipterans such as mosquitos (Lindh *et al*., 2008).

A subset of 27 bacterial ASVs was present in all life stages (Fig 3A), whereas 28 ASVs were shared between adults and eggs and additional 17 ASVs between larvae and pupae. This similarity in taxonomic features indicates the transmission of bacteria across life stages. The larval and egg stages had 118 and 71 unique ASVs, respectively, that were not present in other life stages, indicating a distinct community structure. To assess if and to what degree an inoculation of eggs by adult females might take place, constellations of unique and shared ASVs between FA, EO, WS, and EA samples were calculated in a Venn diagram (Fig 3B). These samples consist of eggs sampled directly from the female abdomen and the emptied female abdomen, a wash of the egg-laying apparatus and the eggs that were directly oviposited afterwards. From this analysis, a profile of shared bacterial groups associated with females and eggs post- and pre-oviposition was derived. The results show that with 22 shared bacterial ASVs between EA and WS it is likely that an inoculation occurs during the oviposition process. Furthermore, the EA has zero shared features with the EO eggs, indicating no similarities before and after oviposition. However, the FA samples shared only three ASVs with EA, indicating that the bacterial community of adult females is not similar to that of freshly oviposited eggs before oviposition starts. This could mean that females accumulate certain bacteria during oviposition to inoculate eggs with a specific community. A similar progression can be derived from the diagrams showing community compositions (Fig 1).

This study aimed at analysing the microbiota on the egg surface and their origin. Our results show that the microbial diversity significantly decreases across the sampling stages and is lowest for the egg stage. This early stage, however, is strongly dominated by members of the *Burkholderiaceae*. The presence of this family of bacteria through all life stages points at a transmission across stages, which becomes especially noticeable between the adult and egg stages during oviposition (EA and WS, Fig 3B). After oviposition, the *Burkholderiaceae* outcompeted other taxa and according to the surface-sterilized eggs, this development seems independent from the adult flies (Fig 1A and C, Fig 2), indicating no active inoculation. To further understand this process in detail, eggs separated from adults right after oviposition should be analysed. Also, different egg surface sterilization methods and their effectiveness should be evaluated.

## Conclusion

The characterization of bacteria associated with the surface of BSF eggs contributes to understanding the important role of microorganisms in interkingdom interaction and the attraction of conspecific gravid females (Zheng *et al*., 2013a). Our results indicate a high abundance of *Burkholderiaceae* in the ES treatment as well as in the EC treatment. Further investigations should be performed to analyse if *Burkholderiaceae* is mainly inside the egg or on its surface. Decrypting this interkingdom interaction could help to manipulate and optimize the process of oviposition via artificial inoculation of a substrate with microbial attractants. This might be a milestone for BSF industrial rearing, as larval loss caused by uncoordinated oviposition could be minimized. Furthermore, by analysing bacterial communities of BSF life stages, this study improves the understanding of the transfer of possible pathogens between life stages and generations.

## Abbreviations

GHG: Greenhouse gas
BSF: Black Soldier Fly
BSFL: Black Soldier Fly larvae
LH: larval haemolymph
GS: Guts from sterilized diet
GU: Guts from a non-sterilized diet
CP: Pupal cell pulp
WS: Wash of egg-laying apparatus
EA: Egg oviposited directly after the wash
EO: egg extracted out of the ovary
FA: Females abdomen after extraction of ovaries
EC: eggs of the fly cage after forced contact with adults
ES: sterilized eggs
DAH: days after hatching

## Supplementary Material

**Fig S1.**
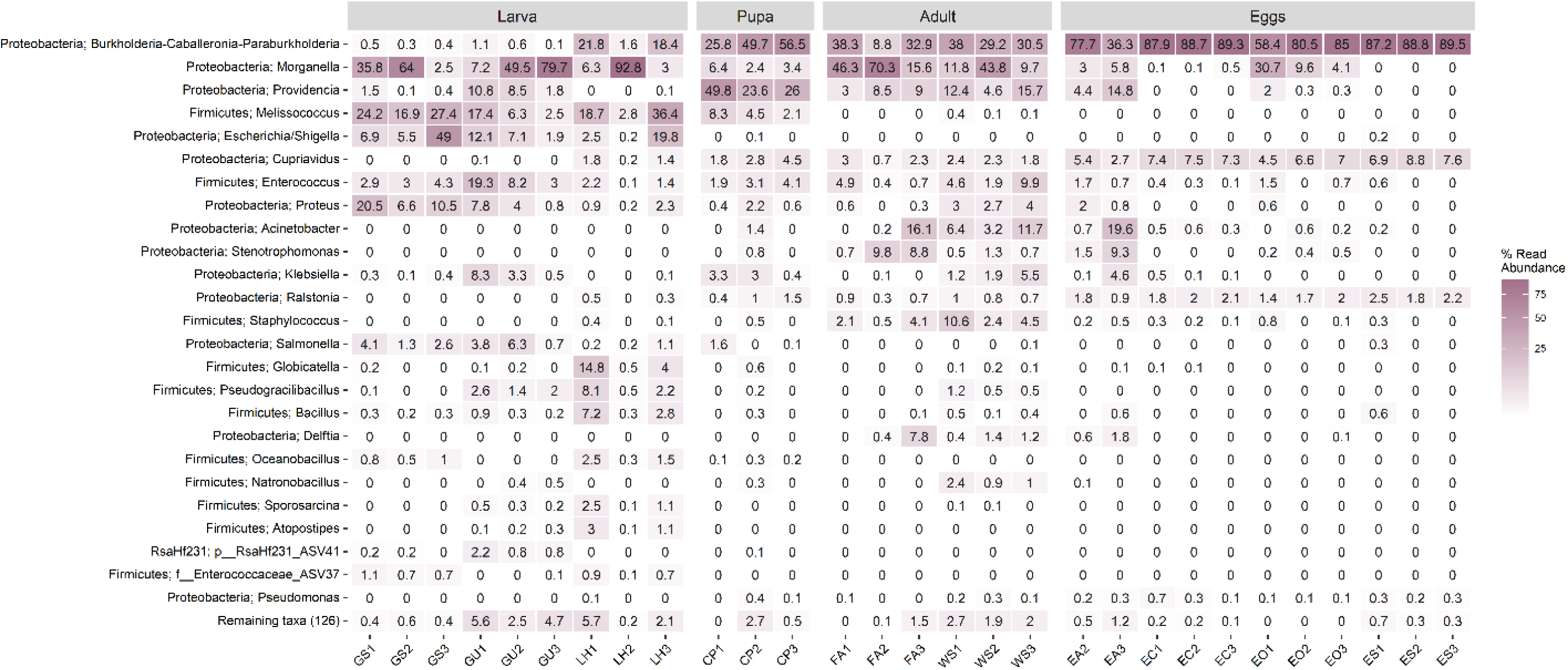
Heatmap of microbial communities in different life stages (A) and tissue samples (B) of the black soldier fly: Larval fed with unsterile (GU) and sterile (GS) feed and the larval haemolymph of GS (LH), the pupal cell pulp (CP), and from the female adults after mating, a wash of the embryo-laying apparatus (WS) and the afterwards placed embryos of the embryo-laying apparatus (EA), embryos collected from the ovary (EO) and the empty female abdomen (FA), embryos collected from a fly cage after forced exposure to adults (EC) and sterilized (ES)

**Fig S2.**
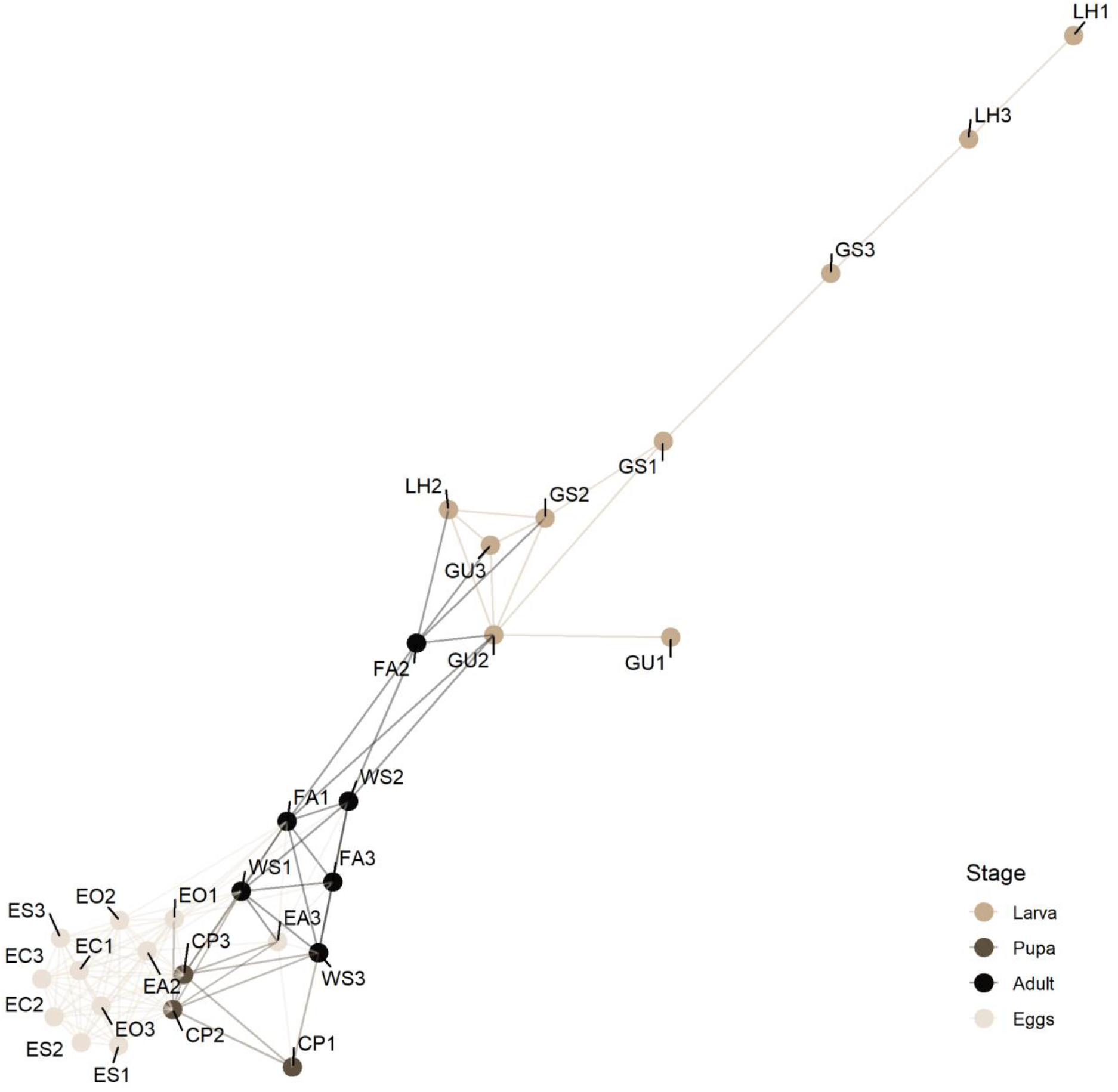
Network analysis of microbial communities in different life stages (A) and tissue samples (B) of the black soldier fly: Larval fed with unsterile (GU) and sterile (GS) feed and the larval haemolymph of GS (LH), the pupal cell pulp (CP), and from the female adults after mating, a wash of the egg-laying apparatus (WS) and the afterwards placed eggs of the egg-laying apparatus (EA), eggs collected from the ovary (EO) and the empty female abdomen (FA), eggs collected from a fly cage after forced exposure to adults (EC) and sterilized (ES)

**Tab S1.** Permanova based on Bray-Curtis analysis in different life stages (A) and tissue samples (B) of the black soldier fly: Larval fed with unsterile (GU) and sterile (GS) feed and the larval haemolymph of GS (LH), the pupal cell pulp (CP), and from the female adults after mating, a wash of the egg-laying apparatus (WS) and the afterwards placed eggs of the egg-laying apparatus (EA), eggs collected from the ovary (EO) and the empty female abdomen (FA), eggs collected from a fly cage after forced exposure to adults (EC) and sterilized (ES). Signif. codes: 0 ‘***’ 0.001 ‘**’ 0.01 ‘*’ 0.05 ‘.’ 0.1 ‘’ 1

|  | Df | SumsOfSqs | MeanSqs | F.Model | R2 | Pr(>F) |  |
| --- | --- | --- | --- | --- | --- | --- | --- |
| ## Stage | 3 | 3.9545 | 1.31815 | 12.676 | 0.60335 | 0.000999 | *** |
| ## Residuals | 25 | 2.5997 | 0.10399 |  | 0.39665 |  |  |
| ## Total | 28 | 6.5541 |  |  | 1.00000 |  |  |

B)
|  | Df | SumsOfSqs | MeanSqs | F.Model | R2 | Pr(>F) |  |
| --- | --- | --- | --- | --- | --- | --- | --- |
| ## Tissue | 4 | 3.6108 | 0.90270 | 7.3607 | 0.55092 | 0.000999 | *** |
| ## Residuals | 24 | 2.9433 | 0.12264 |  | 0.44908 |  |  |
| ## Total | 28 | 6.5541 |  |  | 1.00000 |  |  |

**Tab S2.**
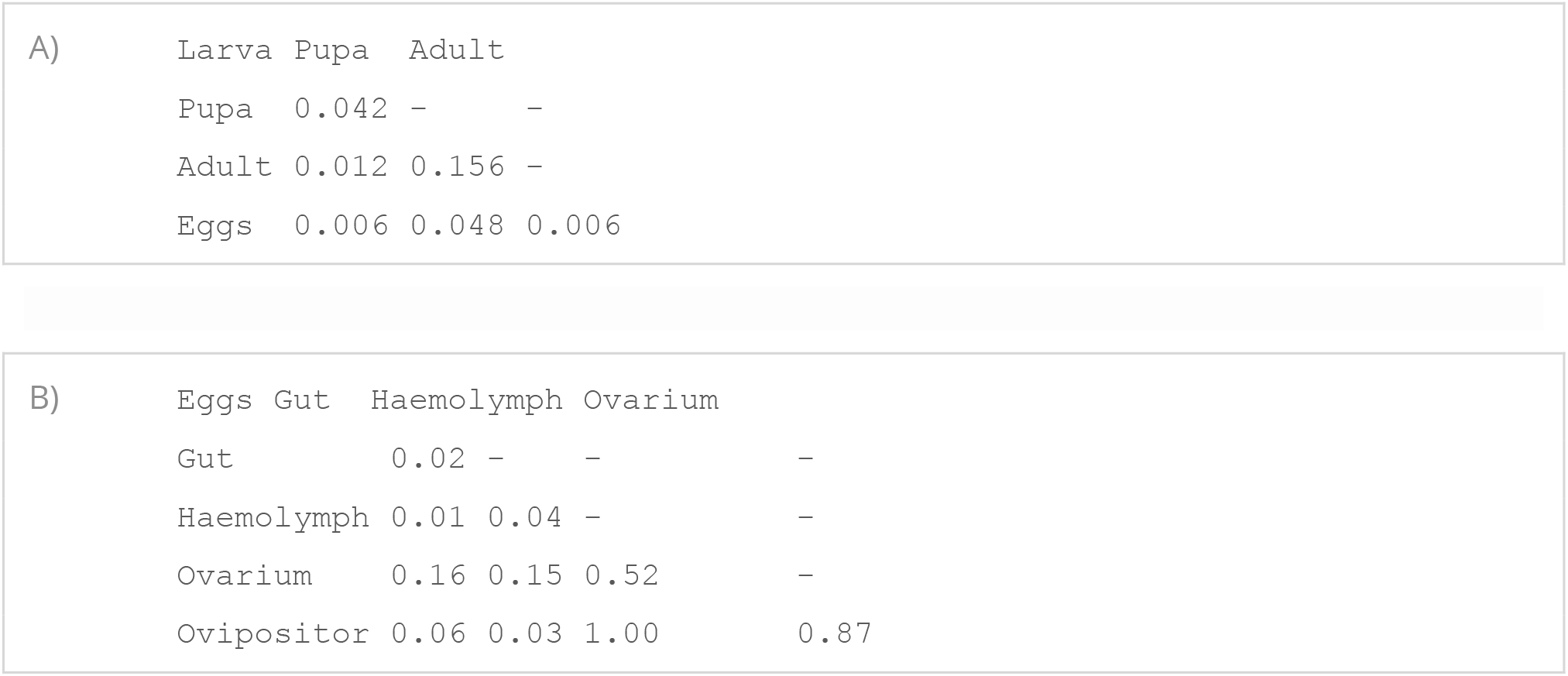
Pairwise Permanova analysis with Bonferroni correction in different life stages (A) and tissue samples (B) of the black soldier fly: Larval fed with unsterile (GU) and sterile (GS) feed and the larval haemolymph of GS (LH), the pupal cell pulp (CP), and from the female adults after mating, a wash of the egg-laying apparatus (WS) and the afterwards placed eggs of the egg-laying apparatus (EA), eggs collected from the ovary (EO) and the empty female abdomen (FA), eggs collected from a fly cage after forced exposure to adults (EC) and sterilized (ES). Signif. codes: 0 ‘***’ 0.001 ‘**’ 0.01 ‘*’ 0.05 0.1 ‘’ 1

**Fig S3.**
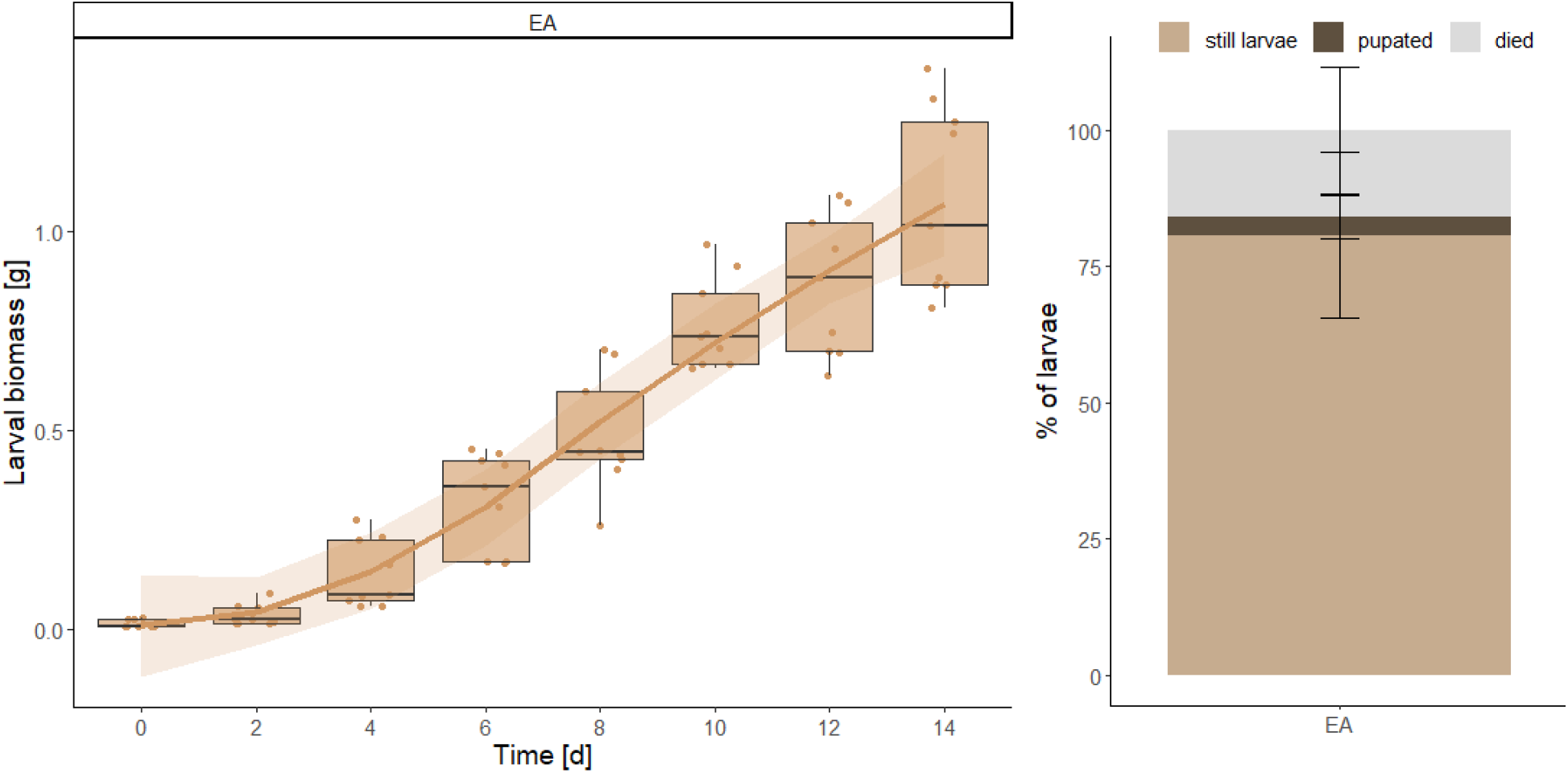
Black soldier fly larvae growing performance, over time, after sampling procedure of forced oviposition of eggs into a sterilized Eppendorf tube (EA) and percentage of pupation and mortality of larvae.

**Fig S4.**
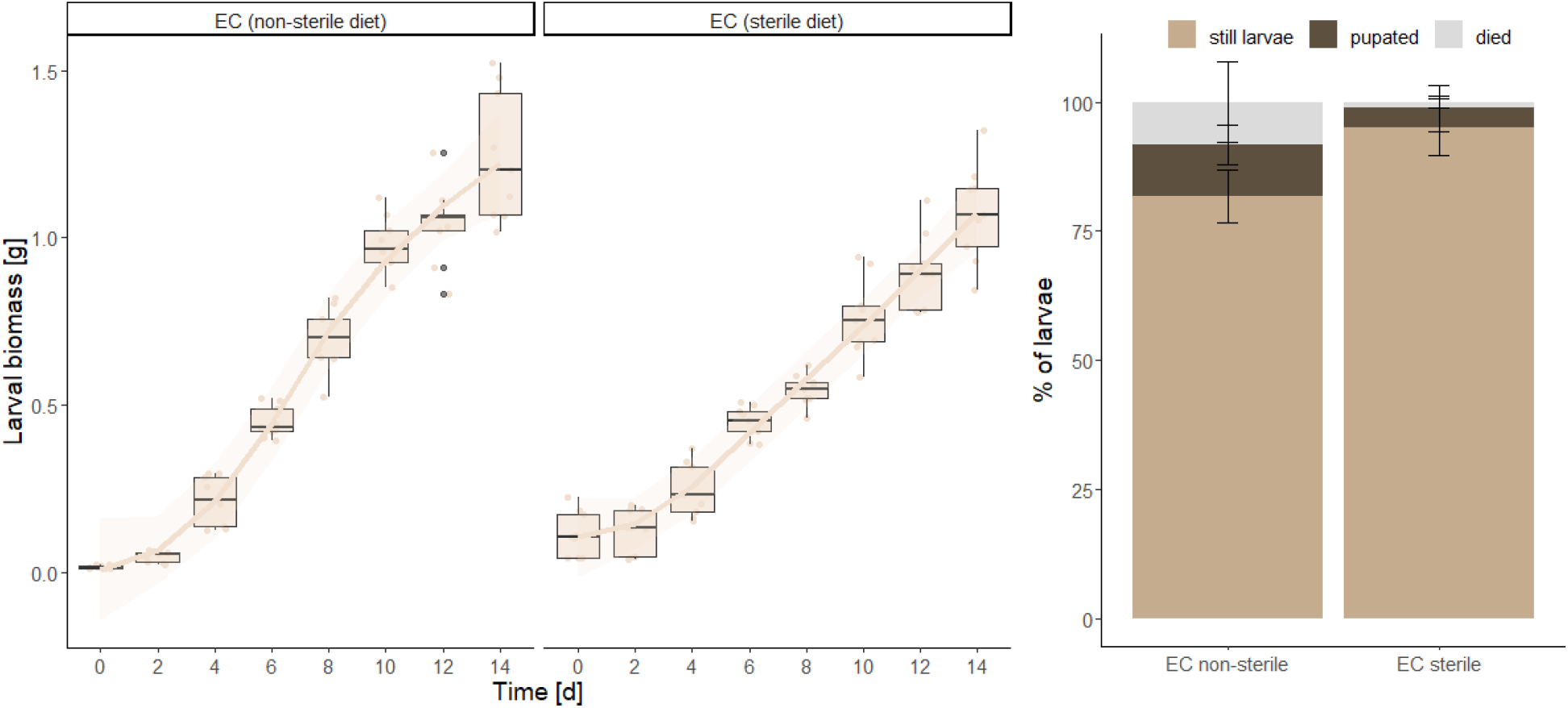
Differences in black sodlier fly growth perfromance reared on a sterilized and non-sterilized feed (chicken feed 40:60 w/v) and percentage of pupation and mortality of larvae.

## Notes

https://tklammsteiner.github.io/eggsurfacemicrobiome

### Competing Interest Statement

The authors have declared no competing interest.

